# Possible environmental and meteorological factors underlying scaly-breasted munia *Lonchura punctulata* (L.) mass mortality events in Southeast Asia’s urban and rural areas in September 2021

**DOI:** 10.1101/2021.09.20.461103

**Authors:** Andri Wibowo

## Abstract

Mass bird mortality is a rare event. This event could be happened and could be a combination of numerous factors. It could be something that’s still completely unknown to us. In September 2021 in SE Asia, there were 2 mass bird mortality events of *Lonchura punctulata*. The first event happens on 9 September in rural area at 08.00 AM and second happens on 14 September at 12.00 PM in urban area. The results show that precipitation combined with the wind gust might be correlated with mass mortality. The results showed precipitation factors have contributed 71.18% (R^2^ = 0.714, P = 0.008) to the mortality and followed by wind gust with 28.81% contribution (R^2^ = 0.41, P = 0.08). The meteorological parameter was not the only factors affecting the mortality events. The landscapes in urban and rural areas have experienced fragmentations. Urban areas have severe fragmentation of vegetation covers with remaining vegetation covers were only 17.4% and patch density indices of 0.29. In contrast rural areas still have higher vegetation cover remnants about 21.12% and patch density indices of 0.93. Then severe meteorological events combined with the fragmented habitats *of L. punctulata* may explain the mass mortality of this bird species.

## Introduction

Mass wildlife mortality events are one of the common events in natural ecosystems. This can be ranged from hundreds of whales stranded on the beach to the massive birds falling from the sky. Those mortality events can be caused by natural causes, anthropogenic causes or combinations of both causes (Calvert et al. 2013, Loss et al. 2015). Among those events, mortality events that were also common recently were the massive mortality of avifauna. Records that date back show that bird mortality has happened since 1800 with large avian mortalities during the migration. This first reported mortality is usually associated with extreme weather events. The second mass mortality event was recorded in 1904 and linked to the snowstorm that killed 1.5 million birds.

In January 2011 in Arkansas, beginning at roughly 11:30 PM, there were estimated up to 5,000 red-winged blackbirds, European starlings, common grackles, and brown-headed cowbirds that fell before midnight. In the next decade, mass bird mortality happened again. This time, the mass mortality events were observed in migratory birds. Thousands of migratory birds are across several states in the Southwest and make this to be the largest mass die-off in recent history. Swallows and flycatchers are among the bird species that are dropping dead in states including New Mexico, Arizona, Colorado, and Nebraska. Bird mortality could be a combination of things. It could be something that’s still completely unknown to us.

Indonesia is one of the countries in Southeast Asia’s region that has high biodiversity including birds. Those bird species are distributed in various ecosystems including in urban areas. One of common bird species in urban and rural settings were *Lonchura punctulata.* Recently 2 mass mortality events of *L. punctulata* have been reported. The first event happens on 9 September in rural area at 08.00 AM and second happens on 14 September at 12.00 PM in an urban area. Current examination on bird carcasses shows negative effects of microbial infections as the causes of mortality. Then this study aims to assess the underlying possible environmental factors of *L. punctulata* mass mortality events in Southeast Asia’s urban and rural areas recently happened in September 2021.

## Methodology

### Study area

The study area was in West Java and Bali provinces (Figure 1), Indonesia. In West Java the location was in urban area of Cirebon city. This city is located near the coast. While in Bali province, the location was in rural Pering village of Gianyar district. This village was also located near the coast and representing rural area.

**Figure 1.**
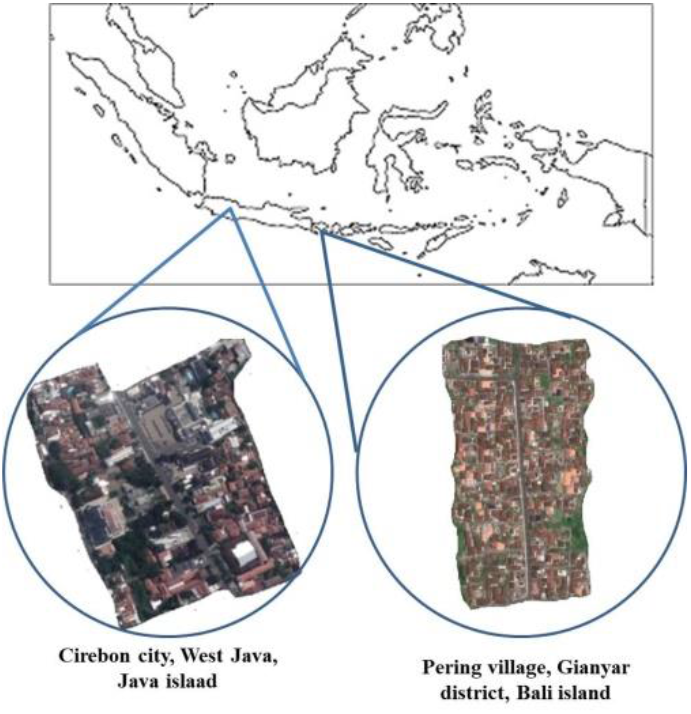
Study area in Cirebon city, West Java province, Java island and rural Pering village of Gianyar district, Bali province, Bali island. Cirebon city represents studied urban area and Pering village represents studied rural area.

### Meteorological analysis

The meteorological data collected including precipitation (inch.) and wind gust (mph) for both study areas. The data was collected for 8 days before bird mortality event happens.

### Fragmentation analysis

Analysis of land cover change was carried out using GIS. The method used followed Samsuri et al. (2018). The measured fragmentation indices were patch area and patch density.

The patch area was formulated:

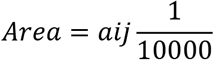

With:

Area = area patch (*m*^2^), divided by 10,000 (convert to be hectare);

a_ij_ = area patch (*m*^2^) patch *ij*

Patch density (Pd) is defined as the number of patch unit on area have 100 Ha as a landscape unit.

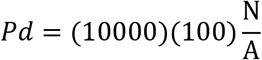

With:

Pd = the number of patch per 100 Ha;

N = the number of forest patch, and

A = area of forest landscape

## Results and Discussions

### Meteorological trends

Meteorological parameters including precipitation and wind gust parameters in event areas were presented in Figure 2. There were similar trends of precipitation parameters both for rural and urban areas. The precipitation trends measured 8 days before bird mortality events were showing an increasing trends. Statistical test confirmed a significant trend of prolonged and escalated rainfalls that might be experienced by the bird flocks. These escalated rainfalls were more significant in rural area (R^2^ = 0.714, P < 0.05, %) rather than in urban area (R^2^ = 0.333, P > 0.05, %) (Table 1). The rainfalls in rural area with averages of 0.0037 inch. (95%CI: 0.00-0.00) were also higher than rainfalls in urban area with averages of 0.0012 inch. (95%CI: 0.00-0.00).

**Table 1.** Statistical test variables (r, P, % contribution) of meteorological parameters (precipitation: inch., wind gust: mph) trends correlated with numbers of days before mortality events

| Area | Parameters | SE | SS | MS | x | (95%CI) | R<sup>2</sup> | P | % Contrib. |
| --- | --- | --- | --- | --- | --- | --- | --- | --- | --- |
| Rural | Precipitation | 0.002 | 0.000 | 0.000 | 0.0037 | (0.00-0.00) | 0.714 | 0.008 | 71.18 |
|  | Wind gust | 3.286 | 45.05 | 45.05 | 10.625 | (12.3- -2.46) | 0.410 | 0.08 | 28.81 |
| Urban | Precipitation | 0.003 | 0.000 | 0.000 | 0.0012 | (0.00-0.00) | 0.333 | 0.134 | 71.18 |
|  | Wind gust | 4.522 | 37.14 | 37.14 | 18.625 | (10.14-46.85) | 0.232 | 0.008 | 28.81 |

**Figure 2.**
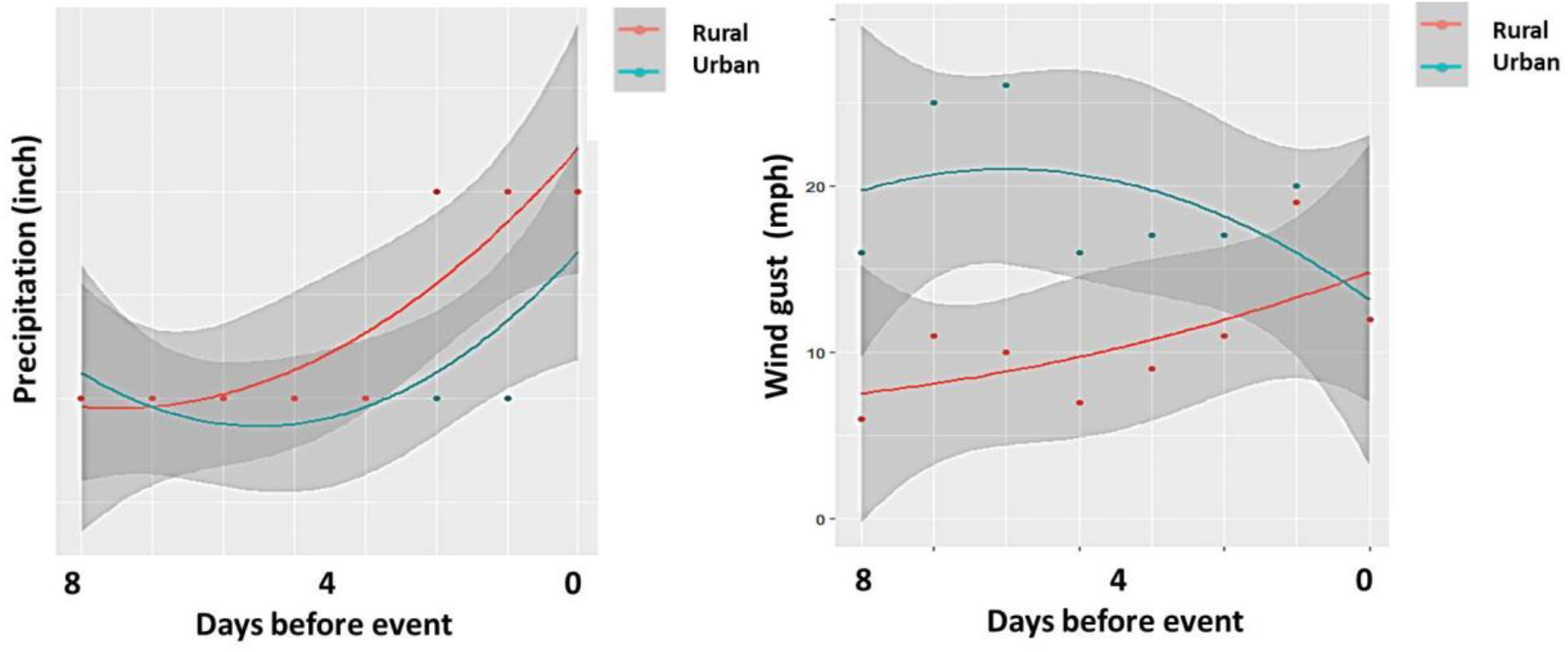
Meteorological parameter trends (95%CI for shaded area) observed from 8 days and 0 day before mortality events. The event was 9 September 2021 for rural area and 14 September 2021 for urban area.

The similar escalating trends for 8 days were also observed for wind gust parameters. Both areas were observed receiving increasing wind gusts. Urban area has experienced higher wind gusts with average of 18.625 mph (95%CI: 10.14-46.85) and average of 10.625 mph (95%CI: 12.3- −2.46) for rural area. While increasing trends of wind gusts were observed more significant in rural area (R^2^ = 0.410, P > 0.05, %) and slightly less significant in urban area (R^2^ = 0.232, P > 0.05, %).

### Consequences on bird mortalities

Bird mortality events were considered as a common event that can occur all the time. In North America, with bird populations of could be as much as 20 billion, almost half die each year due to natural causes. In urban settings, anthropogenic causes can be a significant factor contributing to the bird mass mortalities. This event can be related to loud noises and crashes. A mortality event in Arkansas was reported due to cascading causes. First, legal fireworks set have flushed birds out then forced the birds to fly lower than they normally do, below treetop level, and these birds have very poor night vision and do not typically fly at night. This resulted in bird collisions with cars, trees, buildings, and other stationary objects.

Massive bird mortality in 2020 was also related to human induced natural causes. This mass die-off is happening might be related to several factors including climate change and the ongoing wildfires as possible reasons for the deaths. The wildfires could be to blame, with smoke plumes potentially altering migration routes or increasing toxins inhaled by birds. While, the smoke plume is not the only cause. The smoke plumes are forcing the birds to fly inland and over the desert, which offers little food or water. These events were supported by the fact that some carcasses lacked muscle mass and appeared to have nose-dived into the ground because their faces were damaged. Another cause is might be related to the cold snap in the Mountains West, or a drought in the Southwest of the region that depleted the insect populations that many migratory birds feed on.

The previous aforementioned mass bird mortality events are in agreement with the recent bird mortality event that happened in rural and urban areas. A combination of climatic and meteorological causes (Newton 2017), fragmentation, and possible starving lead to bird mortality can explain the events that happen in rural and urban areas recently. The climatic data including precipitation and wind gust showing significant increasing trends towards especially 4 days before the mortality events. This means that the bird population has been experienced and exposed to climate events in the form of continuous torrential rain that can cause adverse effects. Rain can cause the bird feathers to be wet for a prolonged time and lead to hypothermia (Carr & Lima 2013, Conradie et al. 2020) and even mortality. This is similar to the case reported by Orłowski (2021) that found twenty-eight birds died during intensive rainfall at the end of August and the beginning of September 2001. While during much calmer weather in the same period in 2002, only three birds were killed. Simultaneously, the wind gust factors were also showing increasing trends. A recent study has reported that climatic parameters can contribute to the bird mortalities. McClure reported massive bird mortality following a tornado in 1940 with *Zenaidura macroura* and *Turdus m. migratorius* were the most dominant bird species affected.

Besides massive climatic events that can contribute to bird mortality, another localized meteorological event known as a microburst can be the potential factor. This microburst was in the form of a deadly downward rush of air and sudden wind gust that has uprooted birds roosting from trees. Barometric changes associated with strong thunderstorms can create microbursts that produce a column of quickly sinking air. This condition causes bird to become disoriented and stressed due to the strong weather. Then causing them to rouse during the night and fly into objects. This microburst is a violent burst of sinking air slam straight downward until hitting the ground and spreading out in all directions. The microbursts are very localized and often have high winds capable of knocking over mature trees. Microburst has an adverse impact on wildlife and natural ecosystems. Battles et al. (2017) have recorded that the microburst has removed 4.6% of the canopy of impacted 600 ha area. Within area sizing less than <200 m^2^), 22% of the area damaged by the storm was associated with one 5.2 ha microburst. In this study, a sudden increase in wind gusts has been recorded in both areas. This sudden event might be also contributed to the bird mortalities through microburst mechanisms.

Recently, the weather (Nisbet & Drury 1968) has been reported to become the most important factor influencing bird behavior and limiting bird survival. Bird movement and migration were significantly correlated with high temperatures with presence of high rainfall and wind gust limits the bird movement. This considering that wild birds are much sensitive (Sparks et al. 2002). Increasing precipitation on bird natural habitats has association with bird abundance declines and increasing vulnerability to cowbird parasitism (Rosamond et al. 2020).

Meteorological anomalies were seemingly not the only factors that can contribute to the mass mortalities. In fact, the bird can survive against harsh environmental condition with the requirements that suitable habitats were available. The available habitat in this case is the availability of tree covers indicate and guarantee the foods availability. In fact, the bird habitats in both rural and urban areas were already fragmented with the vegetation covers proportion in comparison to other land covers were less than 20% (Figure 3, Table 2). During the presence of adverse environmental condition including escalating rainfall, high vegetation covers in a certain area is required by birds to have cover for protection and provide food resources. In contrast, fragmentation (McGlinn et al. 2010, Loss et al. 2015) will reduce bird survival especially during escalated adverse meteorological events. While fragmented vegetation covers in urban will increase the exposure of birds to the high speed wind (Laurence & Curran 2008, Damschen et al. 2014) since there is no protection in the form of the tree canopy. Fragmented vegetation covers were also indicating poor food access and resources available for bird to be used as an energy source required to survive during adverse meteorological events.

**Table 2.** Fragmentation indices for vegetation covers in studied rural and urban areas

| Area | Land cover | Landscape proportion | Patch cohesion index | Patch numbers | Patch density |
| --- | --- | --- | --- | --- | --- |
| Rural | 129178 | 21.12% | 0.79 | 573 | 0.94 |
| Urban | 157826 | 17.4% | 0.9 | 268 | 0.29 |

**Figure 3.**
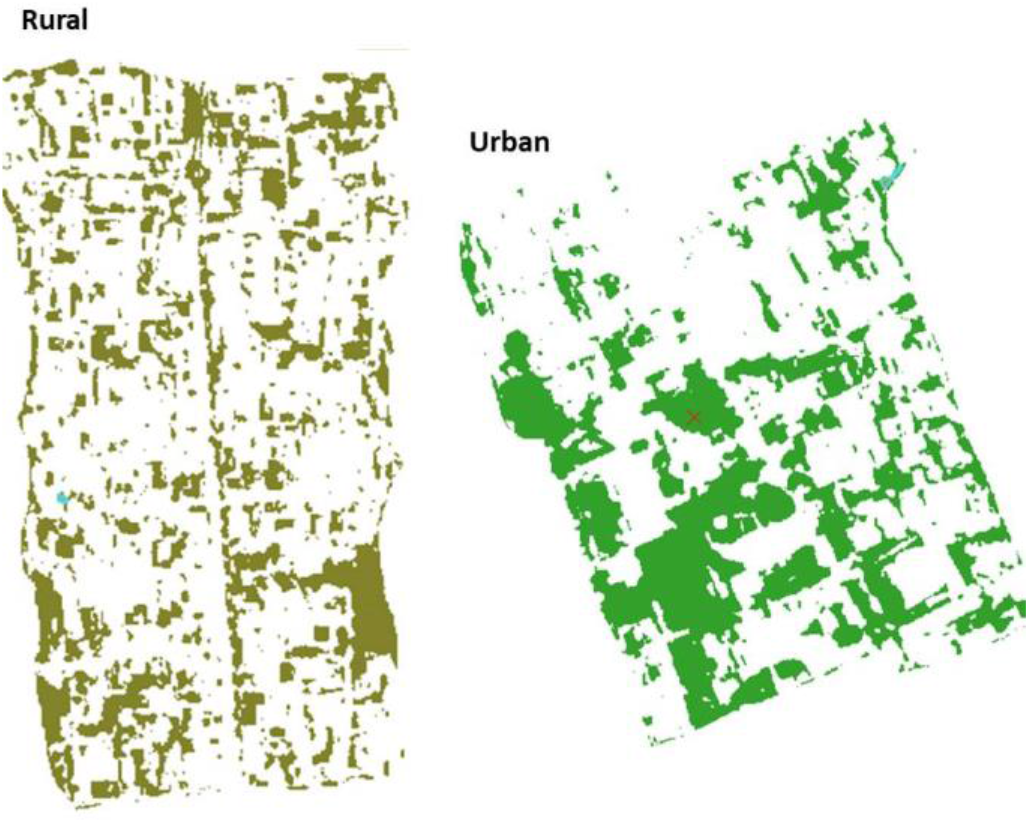
Fragmented vegetation covers in studied rural area (left) and urban area (right).

## Conclusion

Four days before the mass bird mortalities there were increasing trends of precipitation and wind gust. that contribute to mortality. The landscape and bird’s habitat in urban and rural areas were fragmented that have exacerbated the impacts of escalated meteorological events on bird mortality.

## Notes

### Competing Interest Statement

The authors have declared no competing interest.

